# Development of a small-molecule epigenetic regenerative therapy. Subcutaneous administration of alginate formulations with high loads of zebularine and retinoic acid promotes tissue growth, vascularization and innervation and induces extensive epigenetic repatterning

**DOI:** 10.1101/2024.09.18.613177

**Authors:** Paulina Słonimska, Jakub Baczyński-Keller, Rafał Płatek, Milena Deptuła, Maria Dzierżyńska, Justyna Sawicka, Oliwia Król, Paweł Sosnowski, Magdalena Koczkowska, Anna Kostecka, David K. Crossman, Michael R. Crowley, Piotr Sass, Ryszard Tomasz Smoleński, Piotr M. Skowron, Arkadiusz Piotrowski, Michał Pikuła, Sylwia Rodziewicz-Motowidło, Paweł Sachadyn

## Abstract

Recently, zebularine, a small-molecule epigenetic inhibitor and retinoic acid, acting as a transcriptional activator, have been found to induce tissue regeneration. In this study, the pro-regenerative properties of zebularine and retinoic acid were combined with the potential of the alginate carrier to expand its therapeutic possibilities.

Alginate formulations of zebularine and retinoic acid were developed for subcutaneous administration to mice. Hydrophilic zebularine formed a homogenous formulation with extreme drug loadings reaching 240 mg of zebularine per 1 ml of 2% sodium alginate, while hydrophobic retinoic acid, 0.8 mg/ml, dispersed as fine crystals. Cell culture tests exhibited no significant cytotoxicity of the alginate formulations.

Subcutaneous administration of zebularine and retinoic acid in 2% sodium alginate promoted regenerative responses in a mouse model of ear pinna punch wound mice involving the restoration of tissue architecture, nerve and vessel growth, and extensive epigenetic and transcriptional repatterning with no adverse effects observed in the animals. Significant trancriptomic responses to the epigenetic treatment included the induction of epithelium development genes contrasted with the downregulation of muscle development genes on day 7 post-injury. Among the remarkable changes in global gene methylation are those in neurodevelopmental genes. *In vitro* studies showed rapid zebularine but no retinoic acid discharge from the alginate formulations. Live ultrasound imaging demonstrated gradual absorption of the subcutaneously injected alginate formulations, which may explain the *in vivo* activity of retinoic acid following subcutaneous administration.

Effective induction of tissue regeneration together with a high safety profile and of the subcutaneously administered pro-regenerative alginate formulations opens the way to testing further regenerative therapies for hard-to-reach lesions.

## 1. Introduction

Small-molecule drugs are widely used to treat practically all conditions, including infections, cancers, metabolic disorders, and recently even genetic diseases (Laselva et al. 2022), but they remain uncommon in regenerative therapies. Nevertheless, pharmacological induction of tissue regeneration seems a promising direction. Epigenetic mechanisms control processes essential in regenerative response such as cell proliferation, differentiation and pluripotency and therefore attracted researchers’ attention for their role in the regeneration of different organs and tissues, including the liver (Macchi and Sadler 2020, Li and Sun 2024), heart (Jang et al. 2024), muscles (Massenet et al. 2021, Rugowska et al. 2021), skin (Plikus et al. 2015), bones (Wang et al. 2023), and axons (Barker and Tsai 2017, Cheng et al. 2023). Small-molecule epigenetic inhibitors are potential tools that can be used to activate the regeneration process. Our previous research showed that zebularine induced tissue regeneration in an ear-pinna model in mice, thus demonstrating that an epigenetic drug can enhance regenerative abilities (Sass et al. 2019).

Tissue regeneration may take several days or weeks, implying that effective regenerative therapies need extended treatment, achievable with carriers allowing gradual drug release. Hydrogel carriers demonstrate properties functional for regenerative therapies. Hydrogels can be pre-loaded with small-molecule drugs or biologics, easily conform to the application site (Mandal et al. 2020), and gradually release the therapeutic load. The carriers desired for long-term contact with tissues should display excellent biocompatibility. Alginate meets these criteria, making it one of the most extensively tested biomaterials. Alginic acid is an anionic biopolymer of linear chains consisting of β-D-mannuronic acid and its C-5 epimer α-L-guluronic acid linked by 1-4-glycosidic bonds extracted from different brown algae species. It is soluble in water and gels with high porosity (Hamidi et al. 2008). As supported by multiple experiments, alginate is biocompatible (Cattelan et al. 2020). The Food and Drug Administration (FDA) has approved alginate for diverse medical applications, including dental impression materials and wound dressings (Ruvinov and Cohen 2016), as well as oral drugs to treat reflux (Zhao et al. 2020). Alginate is considered non-thrombogenic and non-immunogenic (Ruvinov and Cohen 2016), which may be helpful for the immunoprotection of cell transplants. Alginate entered clinical trials involving implantations of alginate-encapsulated allogenic mesenchymal cells secreting glucagon-like peptide-1 into the brain for the treatment of intracerebral haemorrhage in stroke patients (NCT01298830), intraperitoneal transplantation of alginate encapsulated porcine β-cell islets in patients with type 1 diabetes mellitus (NCT00940173, NCT01736228, NCT01739829), and intracranial implantation of alginate-encapsulated porcine choroid plexus in patients with Parkinson’s disease (NCT01734733). Interestingly, alginate alone may benefit the organism, as reported in the experiment where subcutaneously implanted Alzet pumps with sodium alginate prevented salt-induced hypertension in rats (Moriya et al. 2013).

Retinoic acid, a metabolite of vitamin A, is a transcriptional regulator acting by binding its nuclear receptors, which target retinoic acid response elements in multiple genes (Balmer and Blomhoff 2002, Savory et al. 2014). Retinoic acid, known to be helpful in various skin conditions, has also been approved to treat promyelocytic leukaemia (Yilmaz et al. 2021). Retinoic acid signalling is known to induce regeneration in different tissues in amphibians and mammals (Maden and Hind 2003). Zebularine was tested for its anti-cancer activity as a DNA demethylating agent (Yoo et al. 2004) but has never entered clinical trials. As mentioned above, our previous work demonstrated that zebularine could activate tissue regeneration. What is more, retinoic acid enhanced the regenerative effect (Sass et al. 2019). The treatment involved the administration of several intraperitoneal injections of zebularine and retinoic acid. In the present study, we combined the pro-regenerative activity of the drugs with the potential of the alginate carrier in order to expand the possibilities of therapeutic applications.

## Materials and Methods

### 2.1. Preparation of alginate formulations

Alginic acid sodium salt manufactured from a species of brown algae was purchased from Merck Cat. No. 71238, batch No. BCCD8789). Gorroñogoitia *et al*. presented the characterization of the bioreagent-grade polysaccharide, including its M/G ratio (mannuronic to guluronic acid ratio) and dispersity determined using ^1^H-NMR and gel permeation chromatography (GPC), respectively (Gorroñogoitia et al. 2022). The weight-average molar mass Mw and the number-average molar mass Mn the authors reported were 427 and 186 kDa, respectively, corresponding to the molecular weight distribution (MWD) of 2.3, thus classifying the material to high viscosity alginates. The authors found that mannuronic and guluronic acid contents were 39.6 and 60.4%, respectively, corresponding to a relatively low M/G ratio of 0.7, ranking the tested alginate among those with the highest guluronic content (Gorroñogoitia et al. 2022, Saji et al. 2022). The same authors provided helpful details on the rheological and mechanical properties of the hydrogels prepared from this sodium alginate (Gorroñogoitia et al. 2022).

Alginate carrier was prepared by suspending 20 mg of sodium alginate in 1 ml of water, followed by processing in a bead mill homogenizer (Bead Ruptor Elite, Omni International) using 2 mm porcelain beads at a speed of 4 m/s at room temperature by three 30-s cycles with 10-s breaks. The 2% sodium alginate solutions had a pH of 6.8. To prepare formulations with zebularine, 240 mg of zebularine (TCI, Cat. No. Z0022) was added to 1 ml of 2% sodium alginate and subjected to processing in a bead mill homogenizer by three 30-s cycles with 10-s breaks at 4 m/s using 2 mm porcelain beads. The alginate formulations with retinoic acid were obtained using the same protocol, except 4 mg of all-trans-retinoic acid (TCI, Cat. No. R0064) was added per 1 ml of 2% sodium alginate.

For comparison with non-crosslinked solutions of sodium alginate, the Ca^2+^-crosslinked alginate hydrogels were prepared by the addition of 10 µl of 7 M CaCl_2_ solution to 1 ml of 1 or 2% sodium alginate, followed by bead mill homogenizing performed as detailed above.

### 2.2. In vitro release

Release studies were performed using Phoenix DB-6 Dry Heat Diffusion System (Teledyne Hanson Research, the U.S.A.) equipped with 10 ml vertical diffusion cells with temperature maintained at 37±0.1°C. The system was stirred at a constant speed of 300 rpm. The Spectra/Por Dialysis membrane MWCO: 6-8,000 Da (Spectrum Labs, the U.S.A.) was used to separate the solution/gel from the receptor media. The alginate formulations of zebularine (48 mg per 400 µl) and retinoic acid (0.8 mg per 400 µl) were loaded on the membrane. The diffusion area was 107.5 mm2. All release studies were performed in PBS buffer (pH 7.4). The release experiments were performed in quadruplets. A cumulative release profile of zebularine’s peak area vs time was determined using the UHPLC system, using Phenomenex Kinetex 2.6 µm HILIC 100 Å, 100 x 2.1 mm column in an isocratic mode of 5 mM ammonium formate pH 3.2 in 90% (v/v) acetonitrile in water. The analysis of retinoic acid was performed on the UHPLC system, using Phenomenex Kinetex 2.6 µm C18 100 Å, 100 x 2.1 mm column in gradient mode of 5% to 100% B, where A is 0.1 % of TFA in water and B is 0.1% of TFA in 80% (v/v) acetonitrile in water.

### 2.3. Animals

The experiments were conducted on 8 to 10-week-old females of the BALB/c mouse strain. The animals were purchased from the Tri-City Academic Laboratory Animal Centre, where they were maintained. All animal experiments were performed in compliance with the ARRIVE guidelines. The animal experiment protocols were approved by the Local Ethics Committee for Animal Experimentation in Bydgoszcz (approval no. 51/2020).

### 2.4. Ear pinna punch wound in mice

After the mice were anaesthetized with isoflurane, through-and-through holes of 2 mm diameter were made in the ear pinna centre using a scissor-style ear punch ((Hammacher Solingen; LOT FTC-15/8670/1). Prior to the treatment, the animals were randomized into groups of six. Each group received subcutaneous injections of alginate formulations with zebularine, retinoic acid, their combination, or the carrier alone immediately after wounding (d0) and on day 10 post-injury (d10) in amounts indicated in the legend of Fig. 4. The progress of wound closure was photographed weekly for 6 weeks, followed by the computer-assisted analysis of photographic documentation with ImageJ (Schneider et al. 2012).

### 2.5. Histology

On day 42 post-injury, after the mice were euthanized, the ear pinnae were collected and fixed in 4% paraformaldehyde in 0.01 M phosphate-buffered saline (PBS) pH 7.4 at 4°C. The tissues were then embedded in paraffin, cut into 10 µm sections using a Leica cryostat CM1520, and stained with Masson’s trichrome. Image acquisition was performed with a Leica DM IL LED DMC2900 microscope at 100× magnification.

### 2.6. Immunofluorescence and confocal microscopy

On day 42 post-injury, the animals were euthanized, and both ear pinnae per mouse were collected to 4% paraformaldehyde in 0.01 M phosphate-buffered saline (PBS) pH 7.4 and fixed at 4°C overnight. Next, the ears with the higher degree of wound closure (left or right) from each mouse were selected, transferred to 0.01 M PBS and dissected to outer and inner aspects (closer to the cheek). The outer aspects were used for staining and further analysis. Next, the visible remains of the cartilage layer were gently removed by scrubbing using a small spatula. Then, ears were washed in 0.01 M PBS pH 7.4, 3 times for 5 min and then incubated for 1 h in a blocking buffer comprising of 5% Normal Goat Serum (NGS) in 0.01 M PBS pH 7.4 with 0.5% Triton X-100 at room temperature. Next, the samples were incubated with the primary antibodies diluted 1:300 in 1% NGS in 0.01 M PBS pH 7.4 with 0.1% Triton X-100: monoclonal mouse anti-Tuj1 conjugated with Alexa Fluor-647 (Biolegend, the U.S.A., Cat. No. 801201) to detect neuron-specific class III ß-tubulin in nerve and rat anti CD31/PECAM-1 to detect platelet endothelial cell adhesion molecule-1 as a pan-endothelial marker of vessels (BD Biosciences, U.S.A., Cat. No. 553369). The next day, samples were washed in 0.01 M PBS pH 7.4, 3 times for 5 min. Then, ears were placed on a microscopic glass slide, covered with a mounting medium Vecta Shield Vibrance (Vector Laboratories, the U.S.A., Cat. No. H-1700-10) and a glass coverslip.

Microphotographs of the ear pinnae were captured with the confocal microscope Zeiss LSM800 using ZEN 2.6 Software. Pictures of the wound area were taken using a 10× objective lens, and a 640 nm laser was used for excitation of Alexa Fluor-647 fluorescent dye. Confocal scanning covered the middle 84 μm of the ear’s depth (8 optical slices) to capture all detectable nerve fibres. Photomicrographs were exported as tiff, and a maximum intensity projection of all z-stacks functions was used in ImageJ software to obtain one-plane images. The pictures were calibrated, subjected to uniform background subtraction to minimize the autofluorescence input and changed to 8-bit images. Next, the wound edges were manually drawn to calculate the residual wound area. Then, the regenerated area was determined by subtracting the wound area from the original injury area (3 140 000 µm2). The nerve area was calculated by autothresholding the 8-bit images. Finally, the density was calculated by dividing the nerve area by the regenerated area.

### 2.7. Ultrasound examination

Ultrasound examinations were performed to check the fate of the alginate formulations injected under the skin of mice. The experiments used Vinno6 VET (VINNO Technology, Suzhou, China) and a 21 MHz linear probe. The examinations were conducted on female BALB/c mice aged 8-10 weeks at the beginning of the experiment. During the measurements, the mice were anaesthetized with isoflurane. Measurements were performed immediately after injection of the tested formulations and on days 7, 14, 21, 28, 35 and 42. The experiment involved three mice that received zebularine (48 mg) in 200 µl of 2% sodium alginate and retinoic acid (0.8 mg) in 200 µl of 2% sodium alginate each. Three mice received only 400 µl of alginate alone, and three non-injected mice as naïve controls. Photographs of the ultrasound signal from the area covering the nape and scapulae were exported as tiff, then scaled and transformed to 8-bit images using ImageJ. Next, the intensity auto threshold was applied to convert the images to binary masks and the region of interest, the same for all the analyzed images, was set based on the images from the non-injected naïve mice to cut off the non-specific signal from the ultrasound probe. Five to eight images per mouse per time point were included in the analysis.

### 2.8. RNAseq

Ear pinnae for RNAseq were collected on day 7 post-injury from 6 mice treated with the combination of zebularine and retinoic acid administered in alginate formulations and from 6 control mice receiving the carrier alone, as detailed above (section 2.4). After collection, the samples were immediately placed in liquid nitrogen and stored at -80°C. Total RNA was extracted from 3 mm rings surrounding the initial punch wound using RNeasy Mini kit (Qiagen). The tissues from a pair of ear pinnae from each animal were pooled prior to RNA extraction so that each RNA sample represented one mouse. The quantity of the extracted RNA was determined fluorometrically using the Qubit RNA assay (ThermoFisher Scientific), while the RIN (RNA integrity number) was determined using RNA Screen Tape Analysis on 4150 TapeStation System (Agilent). TruSeq Stranded Total RNA with Ribo-Zero Human/Mouse/Rat kit (Illumina) was used for library construction, followed by 5’ and 3’ adapter ligation. The cDNA libraries were sequenced in paired-end mode on the Illumina platform as a service provided by Macrogen. Sequencing data was converted into FASTQ format with Illumina’s bcl2fastq converter. All samples were trimmed using Trim Galore (version 0.6.7), and these trimmed reads were successfully aligned to Gencode’s GRCm39 Release M28 mouse reference genome using STAR (version 2.7.10a) (Dobin et al. 2013). Transcript assembly and estimation of the relative abundances were performed with HTSeq-count (version 2.0.2). Raw counts were then normalized using DESeq2 (version 1.36.0). The RNAseq data has been deposited in the ArrayExpress database under the accession number E-MTAB-13957.

### 2.9. BSseq

The tissue samples were collected as detailed in the preceding section. Genomic DNA was extracted from 3 mm rings surrounding the initial punch wound using DNeasy Blood and Tissue kit (Qiagen). The tissues from a pair of ear pinnae from each animal were pooled prior to DNA extraction so that each DNA sample represented one mouse. The extracted DNA was quantified spectrophotometrically on the Varioskan Lux (ThermoFisher Scientific), and DIN (DNA integrity number) was determined with Genomic DNA Screen Tape Analysis on 4510 TapeStation System (Agilent). Reduced-representation bisulfite sequencing (RRBS) was performed by the Genomic Core Facility at the University of Alabama at Birmingham using the Ovation RRBS Methyl-Seq kit (Tecan Genomics), followed by sequencing DNA libraries on the NextSeq 500 platform (Illumina). Following Tecan’s Analysis Guide (https://github.com/nugentechnologies/NuMetRRBS), Trim Galore (version 0.6.7) was used to remove residual adapter sequences from the reads. Diversity trimming and filtering with their trimming script were then applied to the trimmed FASTQ files. These clean RRBS sequencing reads were aligned to the UCSC mm39 mouse reference genome using Bismark (version 0.22.1_dev) (Krueger and Andrews 2011). The aligned reads were then sorted using Samtools sort (version 1.15.1), following Bismark alignment, mapping, and sorting, methylKit (version 1.22.0) was used to identify CpG sites, normalize and perform differential methylation analysis. The DMRs were identified with the BIOCONDUCTOR package bsseq using default parameters (Hansen et al. 2012). The BSseq data has been deposited in the ArrayExpress database under the accession number E-MTAB-13955.

### 2.10. Ontological analyses

Genes displaying significant differences in expression (DEGs) with a mean fold change over 2.0 and genes mapped to the DNA regions that showed significant differences in methylation (DMRs) between the treatment and control mice were included in ontological analyses with PANTHER (Thomas et al. 2022). Raw p-values were determined by Fisher’s exact test, following FDR (false discovery rate) correction calculated by the Benjamini-Hochberg procedure to filter out the results with FDR > 0.05.

### 2.11. Cell cultures

The cytotoxicity and the effect on cell viability of alginate formulations were tested on the cultured keratinocytes (HaCaT) and fibroblasts (46BR.1N) treated with the extracts from the alginate formulations. The extracts were prepared by incubating the formulations (1 ml, prepared as detailed above) in 3.3 ml DMEM HG medium (Merck, Cat. No. D6429) for 24 h at 37°C at a surface-to-volume ratio of 3 cm^2^/ml, followed by collection and centrifugation. The cells were seeded into 96-well plates (Corning, Cat. No, 353072), 5000 cells per well, in DMEM HG medium with 10% Fetal Bovine Serum (FBS, (Merck, Cat. No. F9665), and allowed to attach for 24 h in standard conditions (5% CO_2_, 37°C). After removing the medium, cells were treated for 24 h in standard conditions with the extracts from alginate formulations following the LDH (Merck, Cat. No. 11644793001) and XTT (Merck, Cat. No. 11465015001) assays carried out in tetraplicates. The analyses were conducted according to ISO 10993-5:2009 guidelines.

### 2.12. Statistical analysis

Statistical analyses were performed with XLSTAT (Addinsoft). The statistical tests used are indicated in figure legends.

## 3. Results

### 3.1. Preparation of alginate formulations with high loads of zebularine and retinoic acid

In our previous research, to induce the regenerative effects, we used zebularine dissolved in saline and retinoic acid in rape seed oil (Sass et al. 2019). In the present study, we applied the alginate carrier to prepare injectable formulations of both active substances, allowing for gradual release.

Replacing saline with 2% sodium alginate was straightforward, given the excellent zebularine solubility in aqueous solutions reported as 50 mg/ml (Sass et al., 2019). However, the treatment required seven intraperitoneal injections in saline at 500-1000 mg/kg body mass, corresponding to 70-140 mg of collective zebularine dose per mouse weighing 20 g. Bead mill homogenizing facilitated rapid preparation of homogenous injectable formulations containing high zebularine loads (**Fig. 1a**), precisely 48 mg per 200 µl, collectively, 96 mg in two injections. For comparison, trials with Ca^2+^-crosslinked alginate hydrogels were unsuccessful, as such formulations rapidly solidified and were not injectable (**Fig. 1d**). The corresponding load of 240 mg of zebularine per 1 ml of 2% sodium alginate significantly (almost five-fold) exceeded its solubility in aqueous solutions, allowing subcutaneous administration at a reasonable volume of approximately 200 µl per injection site in mice.

**Fig. 1.**
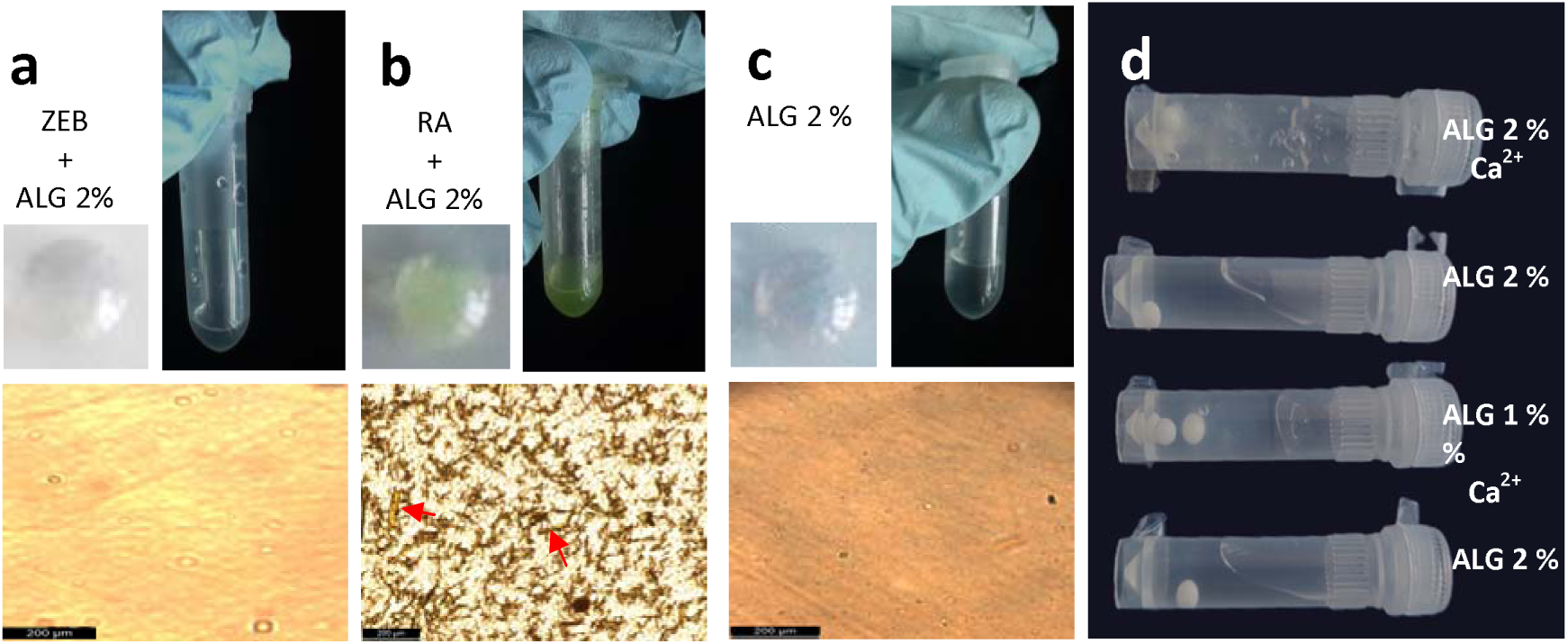
Alginate formulations of zebularine and retinoic acid. **a)** zebularine in 2% sodium alginate (240 mg per 1 ml), a 200 µl drop placed on a plastic dish (top left) in a tube (top right); and observed under a microscope (bottom); **b)** retinoic acid in 2% sodium alginate (4 mg per 1 ml), a 200 µl drop placed on a plastic dish (top left), in a tube (top right); and observed under a microscope (bottom, red arrows indicate tiny crystals of retinoic acid); **c)** for comparison, pure 2% sodium alginate, a 200 µl drop placed on a plastic dish (top left), in a tube (top right); and observed under a microscope (bottom); **d)** comparison of solidified Ca^2+^_-_crosslinked hydrogels with non-crosslinked injectable solutions of sodium alginate that were used to prepare the formulations with zebularine and retinoic acid (the solidified hydrogels do not flow down).

As previously reported, in order to enhance the regenerative effect of zebularine, retinoic aid was administered in rape seed oil in six doses at 8-16 mg/kg, corresponding to a collective dose of 0.96-1.92 mg per 20 g mouse (Sass et al. 2019). Obtaining an injectable alginate formulation of retinoic acid, practically insoluble in water (0.21 µM - 63.1 ng/ml, (Szuts and Harosi 1991), seems more challenging than in the case of zebularine. Hydrophobic molecules can be administered in aqueous carriers if dissolved preliminarily in dimethylsulfoxide (DMSO), but the latter is not desired due to its critical interference with essential life processes (Verheijen et al. 2019). A formulation containing 0.8 mg of retinoic acid in 200 µl of 2% sodium alginate (over sixty thousand-fold exceeding the water solubility of retinoic acid) was obtained using bead mill homogenizing, allowing a colllective dose of 1.6 mg in two injections. The formulation was injectable and uniform under visual inspection (**Fig. 1b**).

### 3.2. Microscopic examination of alginate formulations of zebularine and retinoic acid

Examination with a light microscope did not identify any undissolved reagent residues **(Fig. 1a)**, given that the image resembled that of the formulation of pure 2% sodium alginate (**Fig. 1c**), thus demonstrating the homogeneity of the alginate formulation with zebularine. In contrast, microscopic examination showed fine, uniformly distributed crystals of retinoic acid **(Fig. 1b)**, invisible to the naked eye.

### 3.3. In vitro release of zebularine and retinoic acid from alginate formulations

*In vitro* release was examined in a dry heat testing diffusion system in PBS buffer at pH 7.4 for alginate formulations of zebularine and retinoic acid. The formulation based on 2% sodium alginate gradually released zebularine **(Fig. 2a)**. The process started immediately after placing the formulation in the donor chamber, reaching approximately 33% release within the first 40 min. Over the next 40 min, the cumulative release achieved approximately 50% and approached a plateau of around 70% by 4 h. Then, the process significantly slowed down, reaching the cumulative release of approximately 75% at 24 h and remaining at the same level till the end of the experiment at 76 h. Considering the zebularine half-life of 508 h at 37°C in PBS buffer at pH 7.0 (Cheng et al. 2003), the analysis time was unlikely to impact the results. As expected, retinoic acid, practically insoluble in water (Szuts and Harosi 1991), showed no release from its alginate formulation within the experiment’s timeframe **(Fig. 2b)**.

**Fig. 2.**
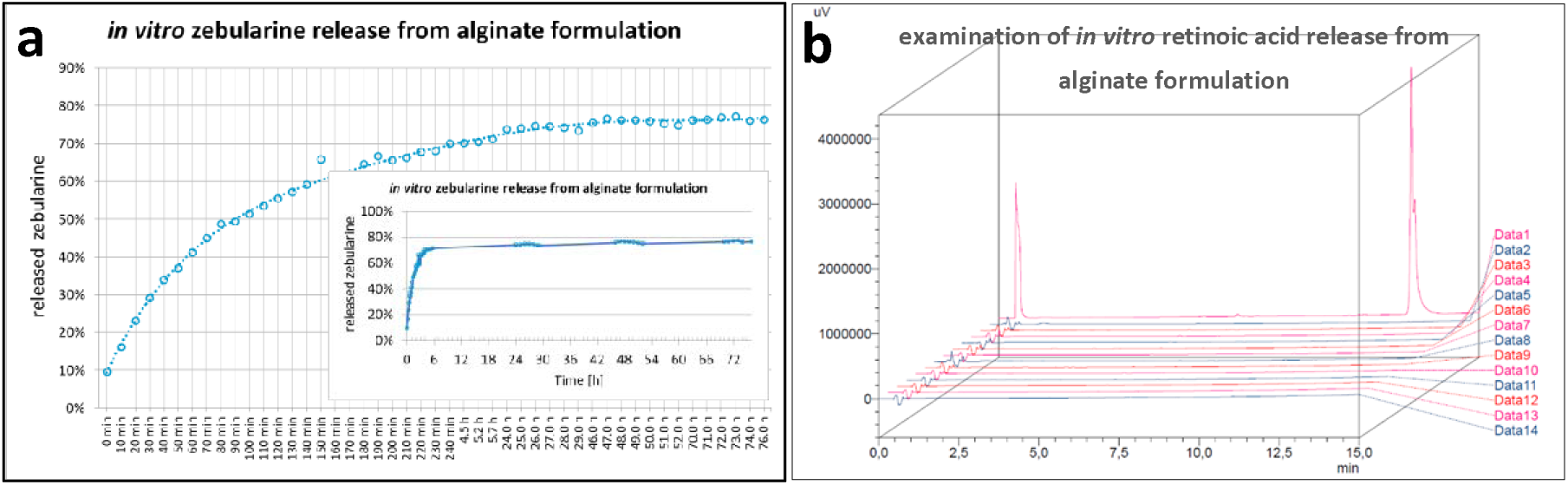
*In vitro* release of zebularine and retinoic acid from alginate formulations. **a)** the percentage of zebularine released to PBS buffer from a formulation containing 48 mg of zebularine per 400 µl of 2% sodium alginate was examined in a dry heat diffusion cell. The main figure presents the results for all time points; in the inset, the data is plotted in the exact time scale to demonstrate the rapid nature of the release **b)** HPLC analysis of retinoic acid release studies from a formulation containing 0.8 mg of retinoic acid per 400 µl of 2% sodium alginate; Data1 – a sample of retinoic acid loaded as a standard, Data2-Data14 – samples collected over time during the release experiment.

**Fig. 3.**
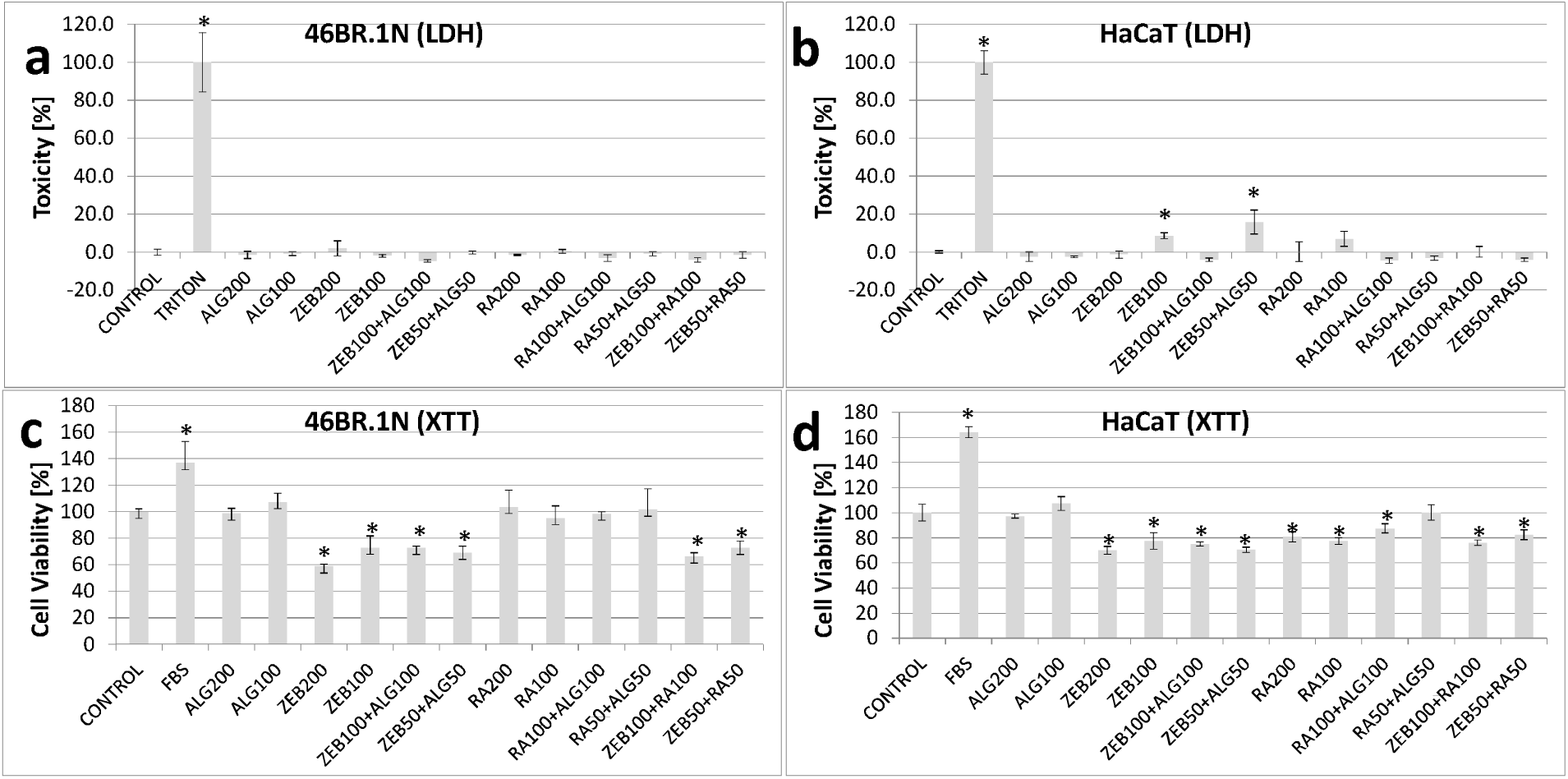
Effects of alginate formulations on cultured fibroblasts (46BR.1N) and keratinocytes (HaCat) assessed with (a, b) cytotoxicity (LDH) and (c,d) cell viability tests (XTT). The cells were exposed to the extracts obtained by incubating alginate formulations in a DMEM HG medium. Each test was performed in 4 replicates; error bars represent the standard deviation (SD). Statistical analysis was carried out using one-way ANOVA followed by *post-hoc* Tukey tests; the significant differences relative to the CONTROL are marked with an asterisk *. The treatments are denoted as follows: **CONTRO**L - cells cultured in DMEM HG without serum; **FBS** - cells cultured in DMEM HG with 10% fetal bovine serum; **TRITON -** cells cultured in DMEM HG with 1% (v/v) Triton X100; **ALG200** - 200 µl of extract from pure 2% sodium alginate formulation **ALG100** - 100 µl of extract from 2% sodium alginate + 100 µl of DMEM HG; **ZEB200** - 200 µl of extract from 2% sodium alginate containing zebularine (240 mg per 1 ml); **ZEB100** - 100 µl of extract from 2% sodium alginate containing zebularine (240 mg per 1 ml) + 100 µl of DMEM HG; **ZEB100+ALG100 -** 100 µl of extract from 2% sodium alginate containing zebularine (240 mg per 1 ml) + 100 µl of extract from 2% sodium alginate; **ZEB50+ALG50** - 50 µl of extract from 2% sodium alginate containing zebularine (240 mg per 1 ml) + 50 µl of extract from 2% sodium alginate + 100 µl of DMEM HG; **RA200** - 200 µl of extract from 2% sodium alginate containing retinoic acid (4.0 mg per 1 ml); **RA100** - 100 µl of extract from 2% sodium alginate containing retinoic acid (4.0 mg per 1 ml) + 100 µl of DMEM HG; **RA100+ALG100** - 100 µl of extract from 2% sodium alginate containing retinoic acid (4.0 mg per 1 ml) + 100 µl of extract from 2% sodium alginate; **RA50+ALG50** - 50 µl of extract from 2% sodium alginate containing retinoic acid (4.0 mg per 1 ml) + 50 µl of extract from 2% sodium alginate + 100 µl of DMEM HG; **ZEB100+RA100** - 100 µl of extract from 2% sodium alginate containing zebularine (240 mg per 1 ml) + 100 µl of extract from 2% sodium alginate containing retinoic acid (4.0 mg per 1 ml); **ZEB50+RA50** - 50 µl of extract from 2% sodium alginate containing zebularine (240 mg per 1 ml) + 50 µl of extract from 2% sodium alginate containing retinoic acid (4.0 mg per 1 ml) + 100 µl of DMEM HG.

### 3.4. Effects of alginate formulations of zebularine and retinoic acid on cultured cells

Previous research in cell culture models demonstrated cytotoxic effects elevated concentrations of zebularine (Sass et al. 2019, Sass et al. 2022) and retinoic acid (Varani et al. 1993, Varani et al. 2007) exert on keratinocytes and fibroblasts. The present study tested the cytotoxic and cell viability effects of zebularine and retinoic acid alginate formulations. For the tests, the cultures of HaCaT keratinocytes and 46BR.1N fibroblasts were treated with extracts obtained by 24-h incubation of cell culture medium at 37°C with alginate formulation containing either zebularine (240 mg per 1 ml) or retinoic acid (4.0 mg per 1 ml) or none. Treating cell cultures with extracts obtained by incubating the tested materials in a culture medium is used to model the interactions of materials with cells and tissues. Such an experimental setup is recommended by ISO 10993-5:2009 guidelines.

No cytotoxic effects were determined in the cultured 46BR.1N fibroblasts (Fig. 3a). The cytotoxic effects in the HaCaT cells were significant only in the case the extracts collected from the alginate formulations with zebularine were diluted with pure culture medium (Fig. 3b, ZEB100, ZEB50+ALG50). In turn, adding the extract from pure 2% sodium alginate counteracted the cytotoxicity (Fig. 3b, ZEB100 vs. ZEB100+ALG100). The observations indicate that zebularine and retinoic acid are released from the alginate formulations, and the extracts from pure 2% sodium alginate, most likely containing dissolved alginate, reduce the cytotoxic activity of zebularine. In addition, zebularine cytotoxicity was neutralized in combined treatment with retinoic acid (Fig. 3b, ZEB50+ALG50 vs. ZEB50+RA50).

Reductions in cell viability by approximately 20-40% were observed in both 46BR.1N fibroblasts (Fig. 3c) and HaCaT keratinocytes (Fig. 3d) after exposure to the extracts collected from the alginate formulations of zebularine and retinoic acid. These effects were significant in the keratinocyte cell cultures for all extracts containing zebularine, retinoic acid, or both, except for that with retinoic acid extract mixed with the extract collected from pure alginate formulations (Fig. 3d, RA50+ALG50). In the fibroblast cell cultures, the decreases were significant only for the extracts obtained from the formulations containing zebularine, either alone or with retinoic acid, but not those with retinoic acid alone (Fig. 3c, RA50+ALG50, RA100+ALG100, RA100, RA200). The effects of zebularine extracts on cultured cells correlate with the observations of zebularine release from alginate formulations *in vitro* (Fig. 2a). While no retinoic acid release was detected *in vitro* (Fig. 2b), the lowered keratinocyte viability (Fig. 3d) indicates that retinoic acid can be freed from its alginate formulations into the culture medium.

In conclusion, the reduced cell viability indicated that alginate formulations released zebularine and retinoic acid into the culture medium. The effect was moderate, considering the extreme zebularine concentration in the alginate formulations. The absence of cytotoxicity demonstrates a remarkable safety profile of the tested alginate formulations. Of note, alginate alone not only had no adverse effects on the cultured cells but even moderated the cytotoxicity of the tested drugs.

### 3.5. Effects of alginate formulation of zebularine and retinoic acid on tissue regeneration

#### 3.5.1. Ear pinna punch wound model in mice

Tests on rodents are widely used in preclinical research to assess the activity of drug candidates. Ear punch wound closure in mice goes beyond a model of cutaneous wound healing. It is an example of tissue regeneration involving not only skin but also cartilage, muscle, vessels, and nerve fibres. Determining the percentage of ear hole closure can be used to identify drugs promoting tissue regeneration (Sosnowski et al. 2022). In the present study, we applied the ear pinna model to test alginate as a carrier for regenerative drugs. The alginate formulations of zebularine and retinoic acid prepared as described above were subcutaneously administered to mice. For reference, analogous treatments were performed with sodium alginate alone, alginate with zebularine alone, and alginate with retinoic acid alone. The treatment was carried out immediately after the injury and repeated on day 10 post-injury. Each tested drug improved ear pinna hole closure compared to the pure sodium alginate control (Fig. 4a,c). Doubling the zebularine dose augmented the regenerative response (Fig. 4b); however, the combined administration of zebularine and retinoic acid had the most significant effect (Fig. 4a). The results were in agreement with those obtained for intraperitoneal administration reported previously (Sass et al. 2019).

The experiment showed that alginate was an effective carrier not only for hydrophilic zebularine readily forming homogenous mixtures in 2% sodium alginate but also retinoic acid, almost insoluble in aqueous media, dispersed in the alginate formulation as fine crystals (Fig. 1b). What is more, both zebularine and retinoic acid in alginate formulations induced pro-regenerative activity after subcutaneous administration. Moreover, the post-mortem sections performed on day 42 post-injury disclosed no remains of the injected material in most (but not all) animals (Fig. 7c). As demonstrated in *in vitro tests,* alginate formulation rapidly discharges zebularine (Fig. 2a). In the case of retinoic acid, subcutaneous absorption of the injected material may explain the release of the drug from the carrier and its activity in the animal’s body (this issue is expanded in the following section 3.6). Although administered in large quantities, the alginate carrier proved safe, as the animals betrayed no adverse effects. No signs of necrosis or irritation at the injection sites confirmed that the formulations were well tolerated under the skin, even though a zebularine concentration of 0.1 mg/ml, way lower than those achieved in the alginate formulations, was cytotoxic, as reported in cell models (Sass et al. 2019).

#### 3.5.2. Histological examination of regenerated ear pinna

Ear pinna hole closure allows preliminary assessment of the regeneration effect, but it is vital to assess not only the extent but also the architecture of restored tissues. A histological examination of the regenerating ear pinnae was performed on day 42 post-injury. The structure of the regrown tissue resembles that of a normal ear pinna with the characteristic presence of the cartilage layer. At the wound edges, developing cartilage can be observed (Fig. 5 b,c,d,e). At the base side, the regrowing ear pinna has a thickness and structure similar to normal tissue, while it widens at the peripheral side. This widening of the peripheral wound edge is less pronounced in the mice treated with zebularine and retinoic acid than in the controls. The peripheral wound end takes a bulb-like structure (Fig. 5c,d,e), except for the mice treated with the combination of zebularine and retinoic acid (Fig. 5b). No cartilage can be distinguished in the bulb-shaped peripheral end in the controls instead, there is an intensive proliferation, and the wound edge is closed by a thick layer of connective tissue (Fig. 5e). In contrast to the controls, skin appendages are found within the regenerating area in mice treated with zebularine and retinoic acid (Fig. 5b,c,d) similar to those in normal tissue (Fig. 5a). In principle, the characteristics of regenerated tissue architecture resemble those observed following zebularine administration in saline reported in the previous research (Sass et al. 2019).

#### 3.5.3. Growth of nerves and vessels within the restored tissues

Ear pinna is supplied by an extensive network of peripheral nerves and blood vessels (Yamazaki et al. 2018). The importance of blood supply in tissue regeneration is well understood, but innervation is also critical, as observed in microsurgical denervation experiments in skin and ear pinna wounds (Buckley et al. 2012, Laverdet et al. 2015, Alapure et al. 2018, Lu et al. 2022). In our experiment, on day 42 post-injury, nerve-specific immunostaining revealed the penetration of the regenerating ear pinna by nerve fibres in both mice treated with zebularine and retinoic acid administered subcutaneously in alginate formulations (Fig. 6b,c,e) and the controls receiving the carrier alone (Fig. 6a,d). Our analysis revealed no significant difference in nerve fibre density in mice receiving the treatment compared to the controls in the circles of 2.4 mm in diameter surrounding the wounds, constituting the area covering the primary injury site (2 mm in diameter) and the peri-injury area affected by the wound healing and remodelling processes (Fig. 6f). Although mice receiving the treatment showed an improved ear hole closure and therefore had more newly grown tissue, the total area of regrowing nerve fibres exhibited no difference compared to the controls. (Fig. 6g). The outcomes indicate that the epigenetic treatment while increasing the area of restored tissue, leads to re-innervation resembling that observed in the controls.

Staining with an endothelial cell-specific antibody revealed dense networks of blood vessels supplying the regenerated tissues (Fig. 6 h,i,j,k,l). As in the case of nerve fibres, there were no significant differences in the densities and the total area of the regrowing vessels between the mice after the epigenetic treatment and the controls (Fig. 6 m,n). Interestingly, the fine nerves and vessels penetrating the regenerating area do not tend to align as the main ones do in the intact ear pinna (Yamazaki et al. 2018).

### 3.6. Biocompatibility – the fate of subcutaneously injected alginate formulations

The post-mortem sections identified no presence of the injected material (Fig. 7c), but whether its elimination was rapid or gradual remained unknown. Live tracking of the alginate formulations injected under the skin of mice using ultrasonography was performed to address this issue. The presence of the injected material was determined by a comparative examination with mice that did not receive injections. The analysis showed gradual decreases in the USG signals from the alginate formulations under the skin, thus suggesting absorption (Fig. 7a,b).

Post-mortem sections indicated no necrosis in the contacting tissues despite huge concentrations of zebularine in the injected formulations (Fig. 7c). The observations correspond with those in cell cultures, where the extracts collected from the formulation of pure 2% sodium alginate mitigated the cytotoxicity of the extracts collected from the formulations containing zebularine (Fig. 3a,b). In addition to no decrease in animal weight observed following the injections (Fig. 7d), the data demonstrate excellent biocompatibility of the tested alginate formulations.

### 3.7.Gene methylation re-patterning and transcriptome changes in regenerating ear pinnae

Genome-wide changes in transcription and DNA methylation following zebularine and retinoic acid treatment were determined using RNAseq and BSseq, respectively. The tissue surrounding wounds (3-mm rings) in ear pinnae was collected 7 days after the injury, and subcutaneous injections of zebularine and retinoic acid in alginate formulations and of the carrier alone. This time point was selected to analyze the critical, early responses to the treatment, early but late enough to record DNA methylation re-patterning. Zebularine-mediated demethylation requires its incorporation into DNA followed by cell proliferation in the growth phase of wound healing (Sass et al. 2019), which will likely begin on day 7 post-injury. The comparison of data from mice receiving treatment compared to those administered with the carrier alone revealed significant changes in gene expression and DNA methylation profiles. Hundreds of differentially expressed genes (DEGs) and differentially methylated regions (DMRs) were identified **(Fig. 8a)**. The mean width of DMR and the mean distance of DMR to the nearest transcription start site were determined as 124 and 6124 bp, respectively. The numbers of hypermethylated regions and downregulated genes decisively prevailed over the hypomethylated and upregulated ones. No remarkable accumulation of DMRs on individual chromosomes was identified **(Fig 8b)**. Most DMRs were located in the intronic and intergenic, while around 34-39% were in the regulatory and exonic regions **(Fig. 8 c,d)**.

**Fig. 4.**
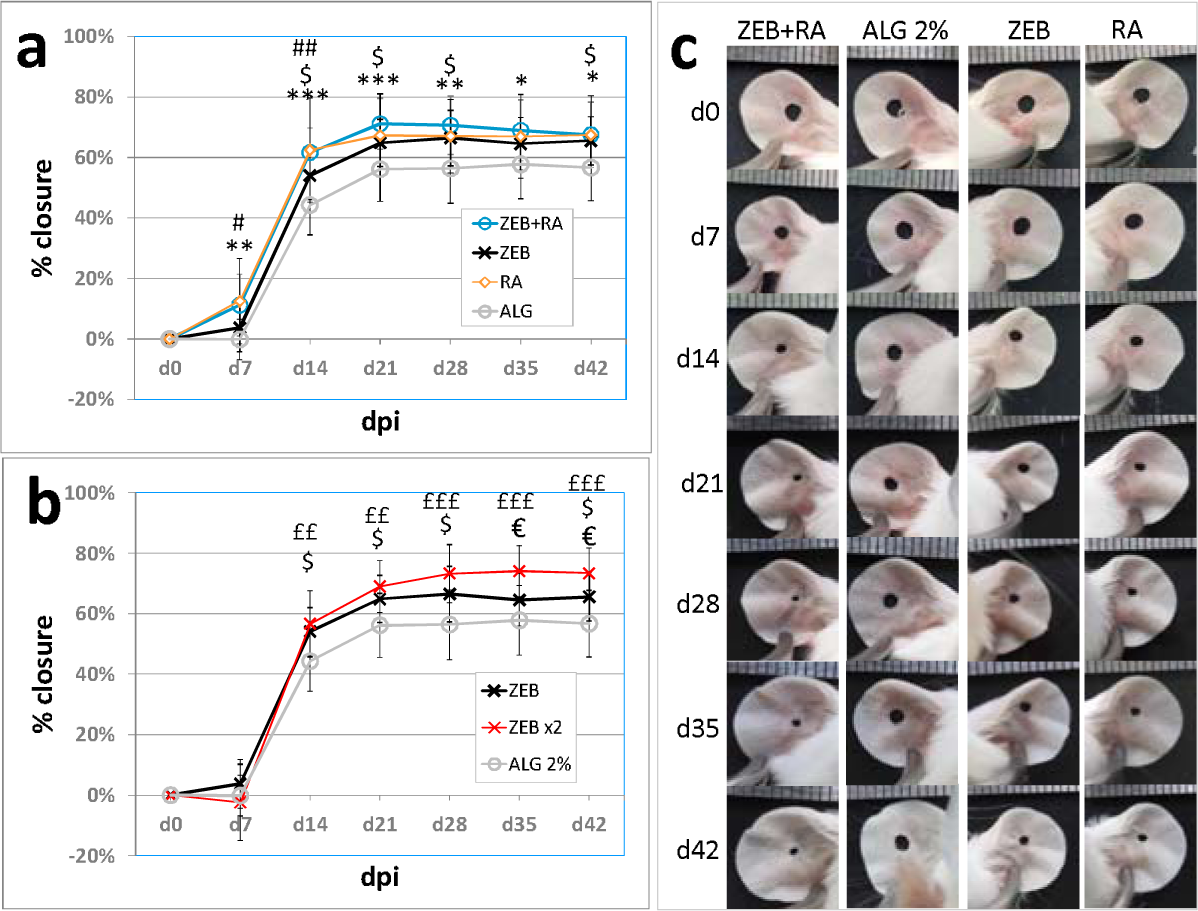
Progress of ear pinna hole closure in mice treated with the formulations of zebularine and retinoic acid in 2% sodium alginate. a, b) mean percentages of ear hole closure for six-mice experimental groups (n=12 ears) receiving subcutaneous injections of alginate formulations on days 0 and 10 post-injury; error bars represent standard deviation. c) representative photographs of ear pinnae. The treatments were designated as follows: ZEB+RA - 48 mg of zebularine in 200 µl of 2% sodium alginate + 0.8 mg of retinoic acid in 200 µl of 2% sodium alginate; ZEB - 48 mg of zebularine in 200 µl of 2% sodium alginate + 200 µl of 2% sodium alginate; RA - 0.8 mg of retinoic in 200 µl of 2% sodium alginate + 200 µl of 2% sodium alginate; ALG - two 200 µl portions of 2% sodium alginate; ZEBx2 - doubled zebularine dose, two 200 µl portions of 2% sodium alginate each containing 48 mg of zebularine each. Statistically significant differences were determined with the two-tailed Mann-Whitney U-test and indicated as follows: with asterisks * for ZEB+RA vs ALG, dollar signs $ for ZEB vs ALG, hashtags # for RA vs ALG, euro signs € for ZEBx2 vs ZEB (doubled vs single dose), and pound signs £ for ZEBx2 vs ALG. Single, double, and triple signs denote p < 0.05, p < 0.001, and p < 0.001, respectively.

**Fig. 5.**
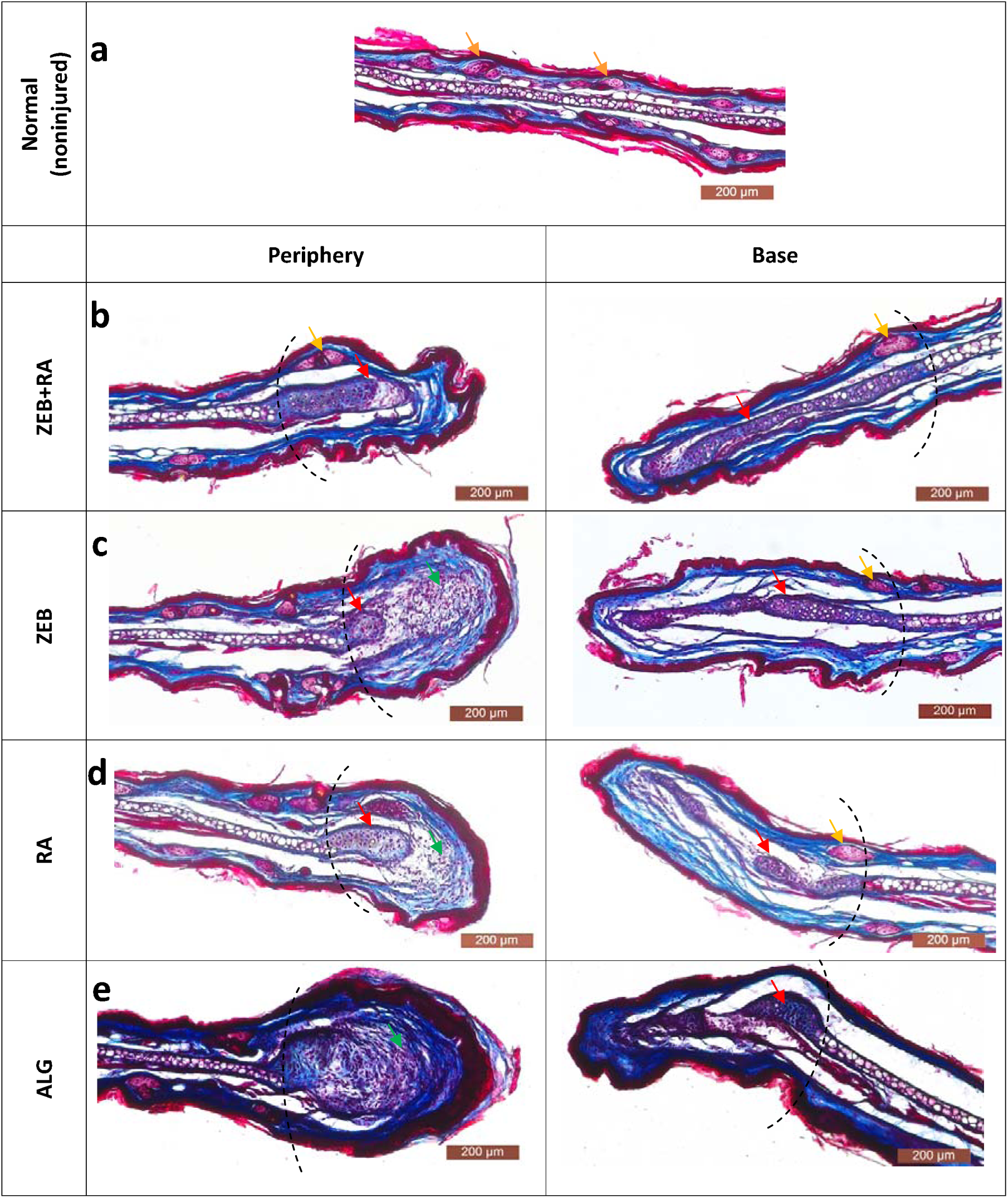
Histological examination of ear pinnae on day 42 post-injury stained with Masson trichrome. The tissue sections were obtained from **a)** noninjured ears (normal tissue); **b)** from injured ear pinnae collected from the mice treated with alginate formulation with zebularine and retinoic acid - **ZEB+RA**; **c**) alginate formulation with zebularine alone- **ZEB**; **d)** alginate formulation with retinoic acid alone - **RA**); and **e)** 2% sodium alginate alone - **ALG**. The dotted lines mark the area of regrown tissue determined based on the comparison of ear hole diameters on days 0 and 42. The red and orange arrows mark the developing cartilage and adnexa, respectively. The green arrows indicate bulb-like structures with cells at the wound margin that were not found in ZEB+RA-treated mice. Staining gives the following results: *epidermis* (keratin) and muscles – red, *dermis* (collagen) – blue, sebaceous glands – purple, nuclei – dark.

**Fig. 6.**
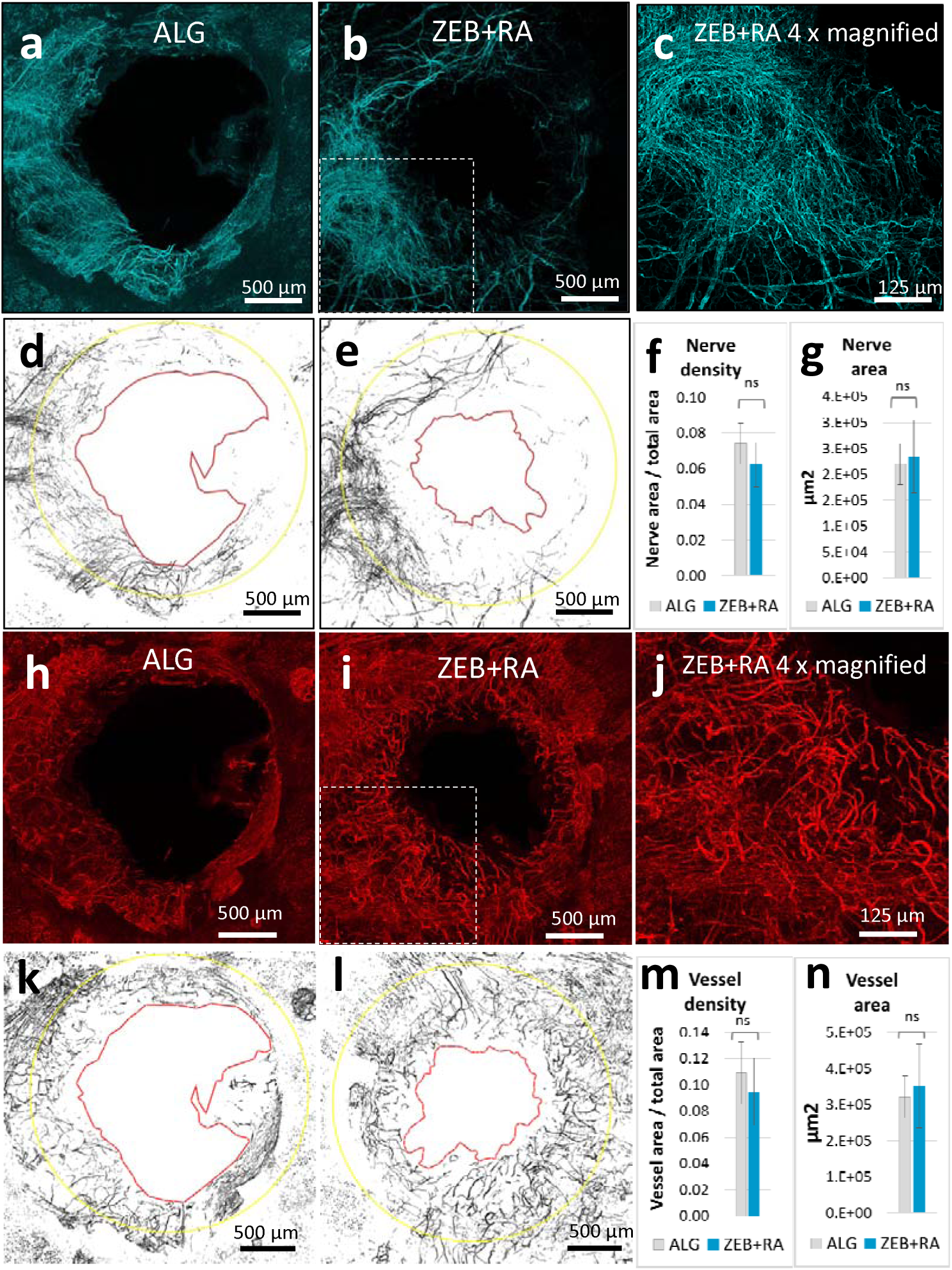
Peripheral nerves and blood vessels growing in regenerating mouse ear pinna on day 42 post-injury. Representative confocal images of the ear pinna outer aspect whole mount stained for neuron-specific anti-β-III-Tubulin (**turquoise pseudocolour**) and endothelium-specific anti-PECAM1 (**red pseudocolour**) antibodies collected from mice treated with **a,h)** pure 2% sodium alginate (**ALG**) and **b,c,i,j)** alginate formulations of zebularine (240 mg per 1 ml) and retinoic acid (0.8 mg per 1 ml) (**ZEB+RA**); binary images of **d,e**) nerve fibres and **k,l**) blood vessels with wound margins on day 42 post-injury marked with red lines and 2.4-mm of diameter yellow circles delineating the area of regrowth; **densities** of **f)** nerve fibres and **m)** blood vessels and **total area** of **g)** nerve fibres an **n)** blood vessels within the regenerating area. Means represent 5 ear pinnae from 3 mice treated with zebularine and retinoic acid (**ZEB+RA**, n = 5) and 6 ear pinnae from 3 mice treated with the carrier alone (**ALG**, n = 6). Error bars represent standard deviation. Statistical analysis was performed using the two-tailed heteroscedastic Student’s t-test.

**Fig. 7.**
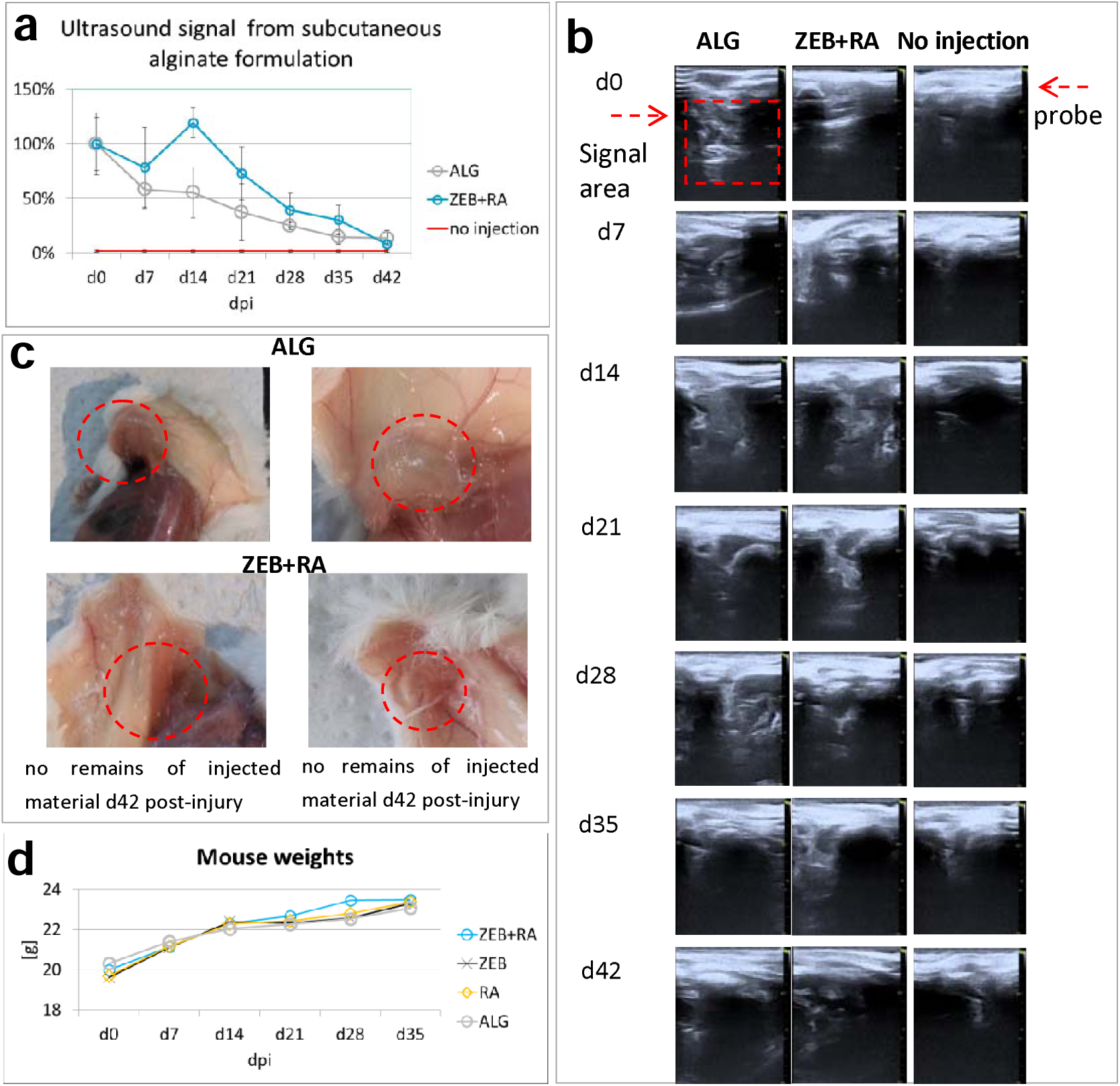
Live-imaging of subcutaneously-injected alginate-based formulations. **a**) ultrasound imaging of subcutaneous alginate formulations measured weekly from the day of injection with 400 µl of 2% sodium alginate denoted as **ALG** and its formulations with zebularine (48 mg in 200 µl of 2% alginate sodium alginate and of retinoic acid (0.8 mg in 200 µl of 2% sodium alginate) denoted as **ZEB+RA**) compared with non-injected mice. Data for each mouse and each time point was determined from 5-8 images. Each treatment was conducted for three mice (n=3); error bars represent the standard error mean (SEM). No significant differences between the groups were determined; **b)** representative ultrasound images; **c)** post-mortem sections on day 42 post-injection; **d)** mouse weights in the course of the treatment with alginate formulations of zebularine and retinoic acid.

**Fig. 8.**
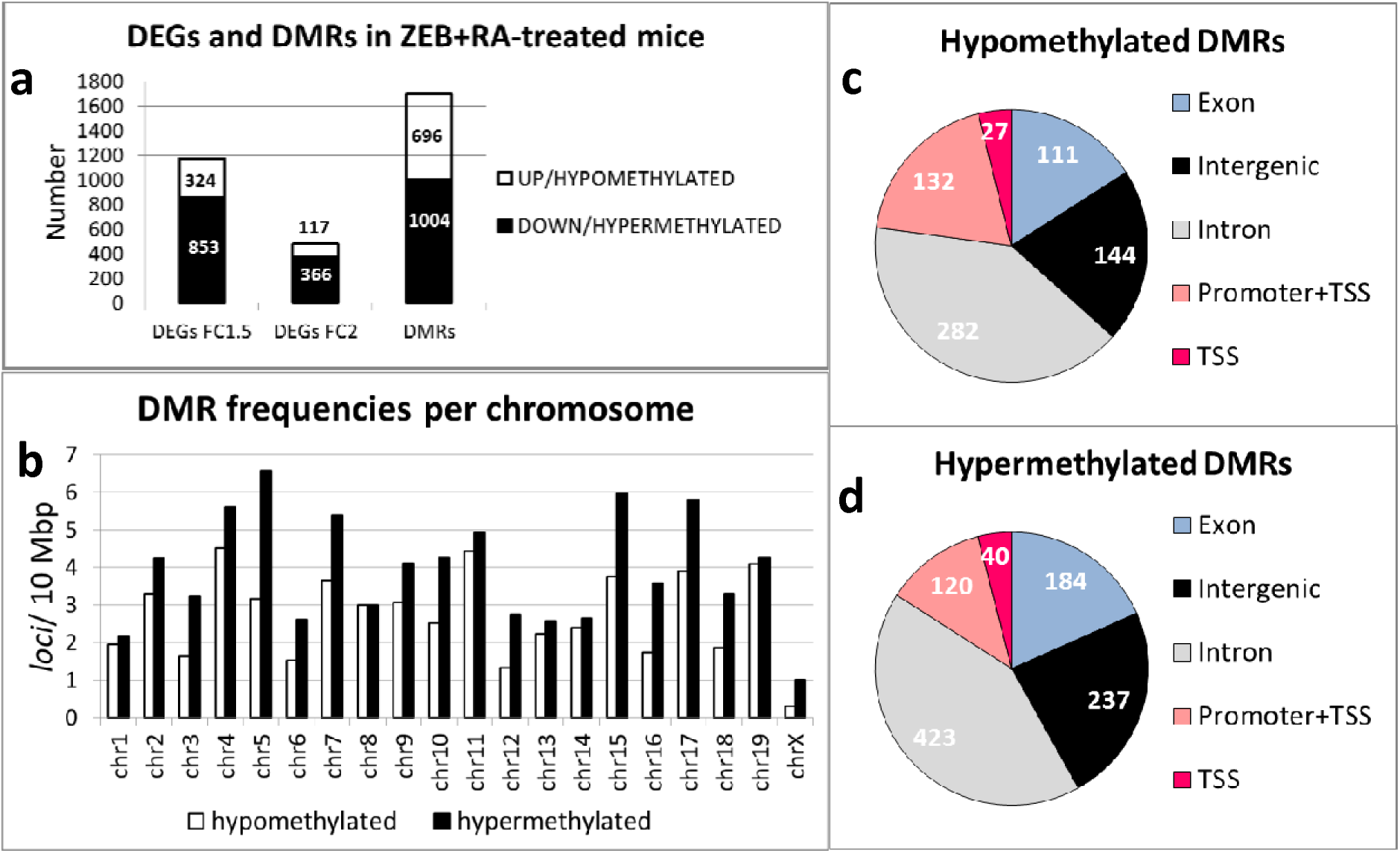
Genome-wide changes in gene expression and DNA methylation in regenerating ear pinnae in response to zebularine and retinoic acid administered in 2% sodium alginate determined using RNAseq and BSseq. Gene expression and DNA methylation were examined in 3-mm tissue rings surrounding 2-mm wounds excised from ear pinnae 7 days after the injury and administration of zebularine and retinoic acid. The tissue samples for RNAseq and BSseq were obtained in independent experiments. Both the RNASeq and BSseq results represent data obtained from 6 mice. **a)** numbers of differentially expressed genes (DEGs) displaying at least a 1.5-fold or a 2-fold change in expression and the numbers of differentially methylated regions (DMRs) in response to zebularine and retinoic acid treatment; **b)** chromosomal distribution of DMRs **c,d)** functional annotation of DMRs.

Ontological analyses were carried out to identify biological processes related to differential gene expression and methylation. The analyses revealed that both differentially expressed genes and those mapped to DMRs were significantly enriched in pathways crucial to regenerative processes **(Tables 1 and 2)**. The pathways identified for the upregulated genes were related to keratinocyte differentiation and epidermis development, while those for the downregulated genes were associated with muscle cell differentiation and muscle tissue development **(Table 1)**. Development, differentiation, and morphogenesis are remarkable for the methylome re-patterning processes, as indicated by the enrichment of genes mapped to DMRs. In particular, "nervous system development" (GO:0007399) is the pathway identified for both the hypo and hypermethylated genes **(Table 2)**. The re-patterning of developmental genes, especially those involved in the nervous system, correlates with our previous findings on the induction of neurodevelopmental genes in ear pinnae regenerating in response to zebularine (Sass et al. 2019). These results correspond with the hypomethylation of developmental genes observed in the models with enhanced regenerative potential (Gornikiewicz et al. 2016, Podolak-Popinigis et al. 2016, Gornikiewicz et al. 2017). The hypomethylated genes, i.e. those potentially activated in the wound margins, showed enrichment in those associated with the Wnt pathway and epithelium development. Of note, "epidermis development" was also the signalling pathway identified for the upregulated genes, whereas "Wnt activation" is characteristic of regeneration (Walczyńska et al. 2023).

**Table 1.** Biological processes associated with the genes enriched in differentially expressed genes (DEGs).

| Upregulated in ZEB+RA-treated mice (3 mm, FC>2, p<0.05) | *raw p-val* | *FDR* |
| --- | --- | --- |
| keratinocyte differentiation (GO:0030216) | 6.02E-10 | 9.35E-06 |
| epidermal cell differentiation (GO:0009913) | 1.55E-09 | 1.20E-05 |
| epidermis development (GO:0008544) | 1.55E-07 | 8.02E-04 |
| skin development (GO:0043588) | 5.75E-07 | 2.23E-03 |
| biological process involved in interspecies interaction between organisms (GO:0044419) | 2.02E-05 | 4.49E-02 |
| Downregulated in ZEB+RA-treated mice (3 mm, FC>2, p<0.05) |  |  |
| muscle structure development (GO:0061061) | 5.08E-10 | 7.90E-06 |
| muscle contraction (GO:0006936) | 3.76E-07 | 8.35E-04 |
| muscle cell differentiation (GO:0042692) | 6.80E-07 | 1.32E-03 |
| muscle tissue development (GO:0060537) | 9.27E-07 | 1.44E-03 |
| regulation of sequestering of calcium ion (GO:0051282) | 7.58E-06 | 1.07E-02 |
| muscle cell development (GO:0055001) | 1.06E-05 | 1.26E-02 |
| monoatomic ion transmembrane transport (GO:0034220) | 3.89E-05 | 3.55E-02 |

FDR correction (false discovery rate) was calculated using the Benjamini-Hochberg procedure.

**Table 2.** Biological processes associated with the genes enriched in the genes mapped to differentially methylated regions (DMRs).

| Hypomethylated in ZEB+RA-treated mice | *raw p-val* | *FDR* |
| --- | --- | --- |
| **nervous system development (GO:0007399)** | 1.15E-07 | 2.55E-04 |
| multicellular organism development (GO:0007275) | 1.02E-07 | 2.65E-04 |
| anatomical structure morphogenesis (GO:0009653) | 8.02E-07 | 9.58E-04 |
| neuron differentiation (GO:0030182) | 1.47E-06 | 1.27E-03 |
| neuron development (GO:0048666) | 9.15E-06 | 4.74E-03 |
| **Wnt signaling pathway (GO:0016055)** | 1.25E-05 | 5.25E-03 |
| embryonic morphogenesis (GO:0048598) | 1.52E-05 | 6.07E-03 |
| nitrogen compound metabolic process (GO:0006807) | 2.49E-05 | 7.91E-03 |
| regulation of cell differentiation (GO:0045595) | 2.77E-05 | 8.26E-03 |
| cell morphogenesis involved in neuron differentiation (GO:0048667) | 2.72E-05 | 8.30E-03 |
| positive regulation of developmental process (GO:0051094) | 4.39E-05 | 1.18E-02 |
| **epithelium development (GO:0060429)** | 5.09E-05 | 1.30E-02 |
| tissue development (GO:0009888) | 1.07E-04 | 2.24E-02 |
| neuron projection morphogenesis (GO:0048812) | 1.08E-04 | 2.25E-02 |
| animal organ development (GO:0048513) | 1.33E-04 | 2.57E-02 |
| programmed cell death (GO:0012501) | 1.65E-04 | 3.05E-02 |
| tissue morphogenesis (GO:0048729) | 1.91E-04 | 3.40E-02 |
| Hypermethylated in ZEB+RA-treated mice |  |  |
| nitrogen compound metabolic process (GO:0006807) | 9.56E-06 | 8.74E-03 |
| **nervous system development (GO:0007399)** | 1.17E-05 | 1.01E-02 |
| carbohydrate catabolic process (GO:0016052) | 2.16E-05 | 1.68E-02 |

FDR correction (false discovery rate) was calculated using the Benjamini-Hochberg procedure.

## 4. Discussion

As reported in our previous research, multiple injections of epigenetic inhibitor zebularine in saline and retinoic acid in oil induced tissue regeneration in ear pinnae in mice (Sass et al. 2019). In the present study, we obtained similar results with two subcutaneous injections of zebularine and retinoic acid in 2% sodium alginate. The regenerative effect manifested in wound closure, tissue architecture restoration, and nerve fibres and blood vessel growth. Significant alterations in methylation and expression status in several hundred genes in the regenerating tissues revealed the extensive scale of epigenetic repattering underpinning the regenerative process. Nervous system development emerged as one of the remarkable pathways associated with the genes differentially methylated and differentially expressed in response to the regenerative treatment. This finding relates to the observed innervation of newly grown tissues.

Alginate proved an effective carrier of both a hydrophilic and a hydrophobic drug, zebularine and retinoic acid, respectively. Applying a bead mill allowed rapid and convenient preparation of injectable formulations with extremely high loads of the active substance (240 mg of zebularine per 1 ml of 2% sodium alginate). The high doses of active compounds in the alginate formulations exerted no adverse effect on the contacting tissues, though zebularine, at elevated concentrations, reduces cell viability and is cytotoxic, as known from previous studies (Sass et al. 2019, Sass et al. 2022). In the cell model, the alginate formulations displayed no significant cytotoxicity but moderately decreased viability of keratinocytes and fibroblasts, thus indicating the release of zebularine and retinoic acid in cell culture conditions. The alginate carrier alone exhibited neither cytotoxicity nor the effect on cell viability.

As shown in *in vitro* experiments, the alginate formulation with zebularine allowed a fast discharge of this hydrophilic compound, which could have been expected, but high amounts of the released substance deserve attention. In contrast, though it forms a homogeneous mixture with 2% sodium alginate solution, highly hydrophobic retinoic acid did not dissolve but formed fine crystals equally dispersed in the alginate formulation. What is conspicuous is that such a formulation did not free retinoic acid *in vitro*. However, both zebularine and retinoic acid activated regenerative responses in mice following subcutaneous injections in alginate formulations. Live ultrasound examinations in the animals showed gradual elimination of subcutaneously injected alginate formulations. The absorption of the carrier most likely explains the regenerative action of retinoic acid administered in the alginate formulation. This observation is important because although alginate is known for its biodegradability, the fate of alginate in mammalian organisms has not been extensively studied. Mammals lack an enzyme specifically digesting alginates (Lee and Mooney 2012). Nevertheless, alginate hydrogel implants were reported as biodegradable (Lansdown and Payne 1994, Novikov et al. 2002, Ueng et al. 2007). On the other hand, Algisyl-LVR, a mixture of alginate with mannitol injected into the left ventricular wall to treat heart failure, displayed long-term stability (Ruvinov and Cohen 2016). The elimination of alginate from the organism may depend on its molecular weight. It has been reported that polymers of alginic acid over 50 kDa are not removed by renal clearance. However, this observation was limited to 24 h following intravenous injection (Al-Shamkhani and Duncan 1995). Our experiments used the alginate with an estimated molecular weight of 100-200 kDa. Gradual fragmentation of the alginate polymers could explain the absorption of subcutaneously administered alginate formulations we observed. The process can be mediated by oxidation, as oxidation was reported to accelerate alginate polymer degradation (Bouhadir et al. 2001, Volpatti et al. 2023).

## 5. Conclusions

Our study demonstrates an application of alginate in epigenetic regenerative therapy used as a small-molecule drug carrier for subcutaneous administration, the route gaining increasing interest from researchers and the pharmaceutical industry (Dubbelboer and Sjögren 2022). Of note is that subcutaneous administration of alginate has rarely been tested, specifically not as a small molecule drug delivery vehicle.

Our observations indicate that both hydrophilic and hydrophobic drugs can be safely administered in alginate formulations in subcutaneous injections following their release. The high load of active substance absorbed by the carrier, enormously exceeding its solubility in aqueous solutions (over sixty thousand-fold in the case of retinoic acid), deserves particular attention. Even if not released *in vitro*, hydrophobic compounds may penetrate tissues following the disintegration and absorption of the injected alginate carrier. This finding is of particular importance for testing compounds with limited solubility in water without using dimethylsulfoxide (DMSO), which is known for critical interference with essential life processes (Verheijen et al. 2019).

Our experiments showed that subcutaneously injected alginate formulations loaded with zebularine and retinoic acid promoted ear pinna regeneration in mice; notably, the process involved the growth of nerves and vessels. Remarkable are alterations in methylation and expression status of several hundred genes observed in the regenerating tissues, specifically the changes in methylation of neurodevelopmental genes. The developed method for drug delivery may help promote regeneration in the case of lesions that are difficult to access directly, like those affecting fine peripheral nerve fibres or capillary blood vessels needed for tissue innervation and vascularization.

## Acknowledgements

This study was supported by the grant "BIONANOVA" of the National Centre for Research and Development of Poland No. TECHMATSTRATEG2/410747/11/NCBR/2019. The funding source had no role in the writing of the manuscript or in the decision to submit it for publication.

We are grateful to the Foster Foundation for their generous support in purchasing the Zeiss LSM 800 confocal la The financial support to maintenance of research facilities used in these studies from Gdańsk University of Technology by the DEC--2/2021/IDUB/V.6/Si grant under the SILICIUM SUPPORTING CORE R&D FACILITIES – “Excellence Initiative - Research University” program is gratefully acknowledged ser scanning microscope we used in our experiments.

The financial support for maintenance of research facilities used in these studies from the Gdańsk University of Technology by the DEC-2/2021/IDUB/V.6/Si grant under the SILICIUM SUPPORTING CORE R&D FACILITIES – “Excellence Initiative - Research University” program is gratefully acknowledged.

We want to acknowledge the support in animal handling from the personnel of the Tri-City Academic Laboratory Animal Centre, the Medical University of Gdańsk, in particular Grażyna Peszyńska DVM, Anna Żyłko PhD, and Agnieszka Jakubiak MSc Eng.

## Author contributions

**P. Słonimska** - Conceptualization, Data curation, Formal analysis, Investigation, Visualization, Methodology, writing original draft, writing - review & editing; **J. Baczyński-Keller** – Investigation, Methodology, Formal analysis, Visualisation; **R. Płatek** - Investigation, Methodology, Formal analysis, Visualization; Writing original draft **M. Deptuła** - Investigation, Methodology, Formal analysis; **M. Dzierżyńska** - Investigation, Methodology, Formal analysis; **J. Sawicka** - Investigation, Methodology, Formal analysis; **O. Król** – Methodology; **P. Sosnowski** – Investigation, Methodology; **M. Koczkowska** - Data curation, Investigation, Methodology, Formal analysis; Writing - review & editing **A. Kostecka -** Data curation, Investigation, Methodology, Formal analysis**, D.K. Crossman -** Investigation, Methodology, Formal analysis**; M.R. Crowley** Investigation, Methodology**, P. Sass** – Methodology; **R.T. Smoleński** – Supervision; **P.M. Skowron** – Funding acquisition, Conceptualisation; **A. Piotrowski** - Investigation, Methodology, Formal analysis; Funding acquisition; **M. Pikuła** – Conceptualisation, Formal analysis, Funding acquisition, Supervision; **S. Rodziewicz-Motowidło** - Conceptualisation, Formal analysis, Funding acquisition; Supervision; **P. Sachadyn** – Conceptualisation, Formal analysis, Funding acquisition, Supervision, Data curation, Investigation, Writing - original draft, and Writing - review & editing, Visualization.

---

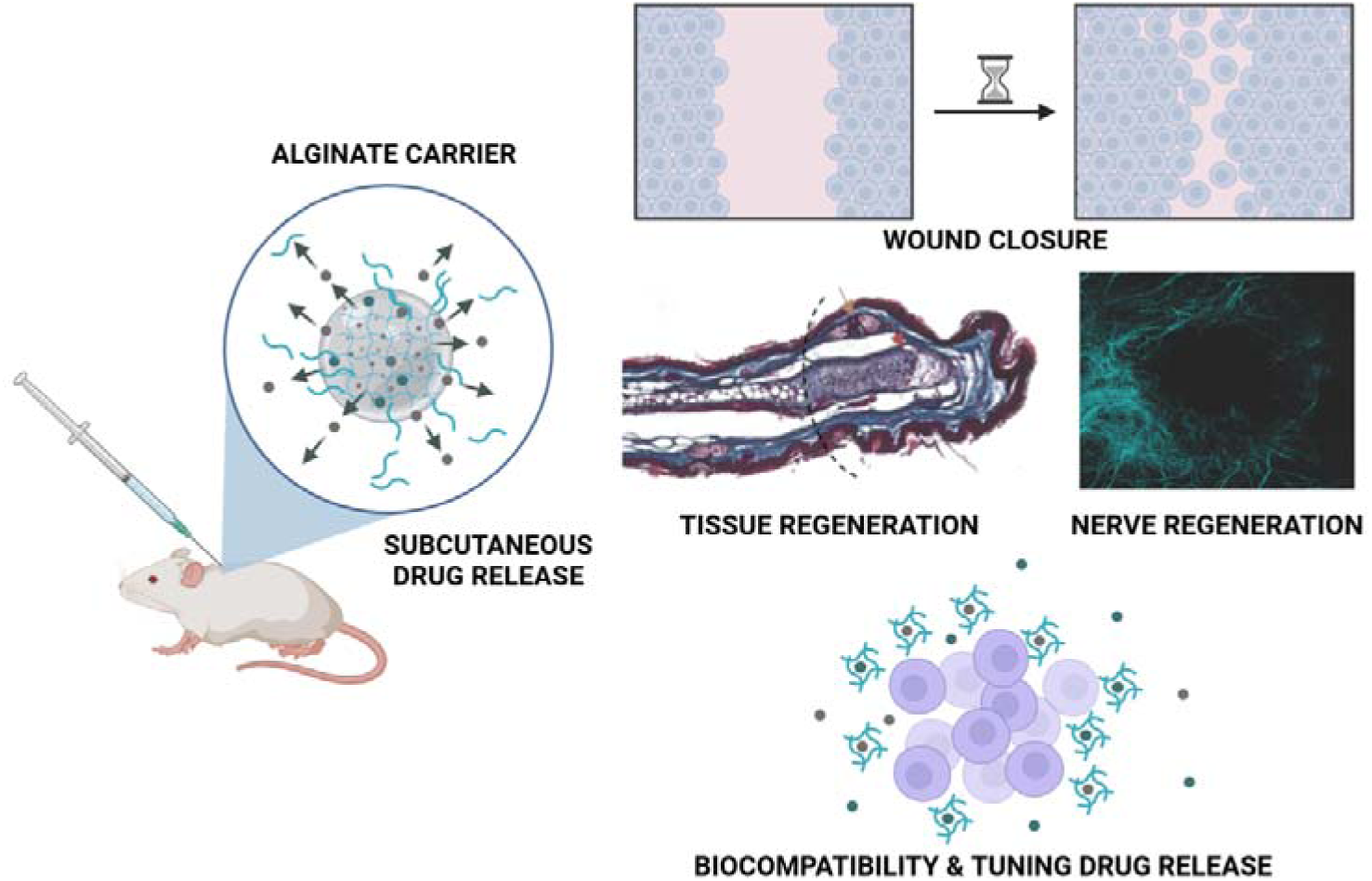

